# Simplest Model of Nervous System. IV. General Solution

**DOI:** 10.1101/2024.07.10.603010

**Authors:** Anton V. Sinitskiy

## Abstract

In this paper, we extend our previous work on a simplified model of the nervous system by solving the general optimization problem for the evolutionary cost of the nervous system. This optimization takes into account constraints on the scales of membrane potential kinetics and sensory response function to ensure finite, biologically plausible solutions. Our analysis reduces the variational problem to a system of two integro-differential equations, which we solve asymptotically using series expansions. This study confirms the emergence of sharp finite neuronal spikes and robust sensory and motor response functions as evolutionarily optimal solutions. We note that, in principle, alternative evolutionary solutions with different biophysical interpretations might exist for this optimization problem. This work provides a rigorous mathematical framework bridging evolutionary optimization with the fundamental properties of nervous systems.

## Introduction

In our previous research,^1-5^ we introduced a novel, highly simplified model of the nervous system, inspired by a hypothetical origin scenario, that effectively captures critical properties of the nervous system. For the formulation of the problem, mathematical definitions, contextual discussion, and literature review, we refer readers to our earlier work.^1,2,5^ Evolutionary optimization in this model is represented as the minimization of the functional *I* defined as:

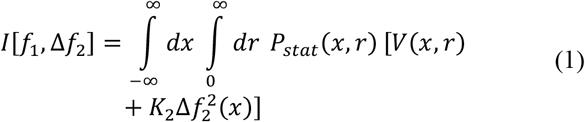

under functions *f*_1_(*x, r*) and Δ*f*_2_(*x*), where:

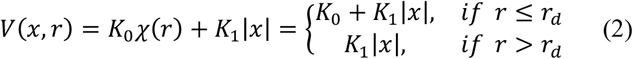

and *P*_*stat*_ (*x, r*) is the stationary probability density, which can be found from the Fokker-Planck-Kolmogorov equation:^6^

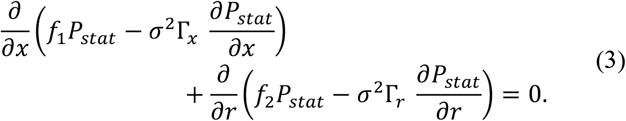

In these equations, *K*_0_, *K*_1_, *K*_2_, *r*_*d*_, *σ*^2^Γ_*x*_, and *σ*^2^Γ_*r*_ are positive constants, *f*_2_(*x, r*) = *f*_*pred*_(*r*) + Δ*f*_2_(*x*), and *f*_*pred*_ (*r*) is a known function. Note that by definition, *I* ≥ 0. For definitions of all these variables and functions, refer to our previous work.^1,2,5^

The stationary probability density *P*_*stat*_ (*x, r*) is normalized to unity:

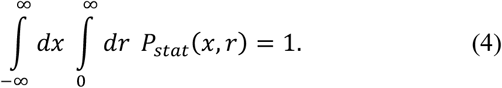

In previous work, we performed a partial optimization of *I* as the functional of only Δ*f*_2_(*x*), assuming that the function *f*_1_(*x, r*) was known. In this work, we lift this restriction and perform a complete optimization of *I* as a functional of both Δ*f*_2_(*x*) and *f*_1_(*x, r*). This generalization requires introducing a restriction on the scale of *f*_1_(*x, r*), which we suggest choosing in the following form:

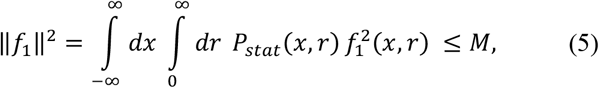

where *M* is some large number. The biophysical interpretation of this constraint is as follows. Material mechanisms responsible for the sensory role of a neuron, as well as the return of its membrane potential to the resting state, operate on a finite scale. For example, for *f*_1_(*x, r*) previously used in our numerical simulations, *f*_1_(*x, r*) = −*ax* + *C* exp(−λ*r*), the constants *a* and *C* may be very large, but not infinitely large, being limited by the conductance of ion channels, finite timescales of molecular diffusion, etc. Given this specific functional form of *f*_1_(*x, r*), we could impose the restrictions *a* ≤ *a*_*max*_ and *C* ≤ *C*_*max*_. However, these restrictions do not apply to the general functional form of *f*_1_(*x, r*). We believe restriction (5) may be the best generalization of these conditions to the general case.

### Transformation of the variational problem to integro-differential equations

The Lagrangian for minimizing *I* from equation (1) under restrictions (3)-(5) can be written as

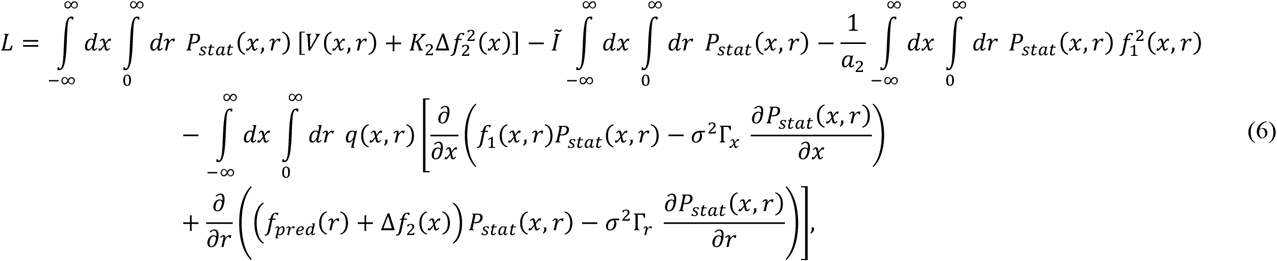

where *Ĩ*, 1/*a*_2_ and *q*(*x, r*) are Lagrange multipliers. We choose the second of them in the form of 1/*a*_2_, because later we will find an analogy between |*a*_2_| and *a*^2^, where *a* is a large parameter from the derivations in paper III.^5^ However, we are unsure about the sign of *a*_2_, so we cannot simply write it as *a*^2^ or −*a*^2^.

Now, in the last term, let us perform integration by parts twice, under the assumptions that

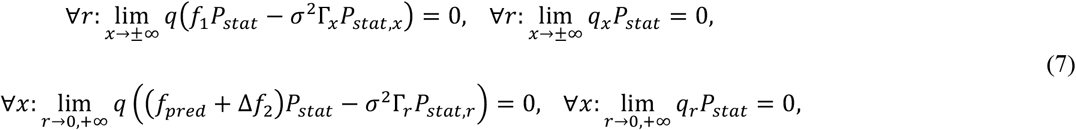

where here and below we use subscripts *x* or *r* to denote the corresponding partial derviatives. This integration leads to the following alternative expression for *L*:

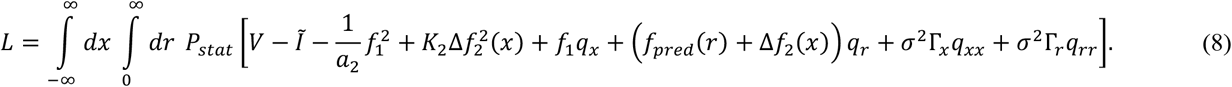

Variation of the Lagrangian with respect to *Ĩ*, 1/*a*_2_ and *q*(*x, r*) reproduces restrictions (4), (5) and (3), respectively, as seen from the Lagrangian given by equation (6). The remaining variations will be performed with the use of the Lagrangian given by equation (8). Variation with respect to *P*_*stat*_ (*x, r*) yields:

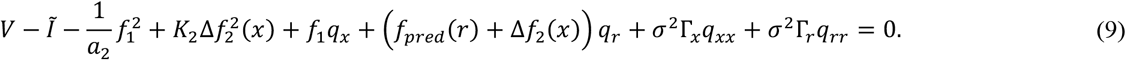

Variation with respect to *f*_1_(*x, r*) results in

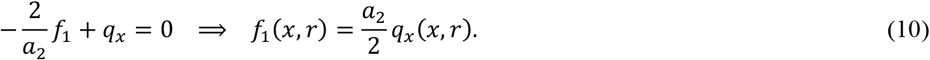

Variation with respect to Δ*f*_2_(*x*) leads to

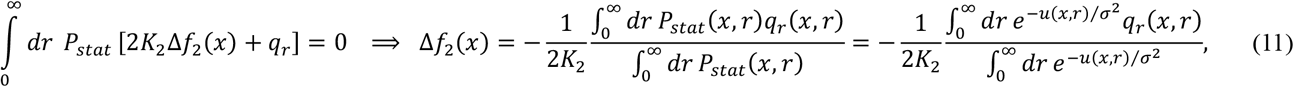

where *u*(*x, r*) is defined from *P*_*stat*_(*x, r*) by equation:

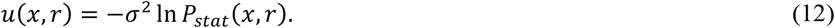

Plugging these expressions for *f*_1_(*x, r*) and Δ*f*_2_(*x*) in terms of *q*(*x, r*) into equation (9), we get a second-order partial integro-differential equation for *q*(*x, r*). Since this equation, besides known constants and functions, contains *u*(*x, r*), it should be solved together with the Fokker-Planck-Kolmogorov equation (3), rewritten in terms of *u*(*x, r*) and with the new expressions for the drift terms in terms of *q*:

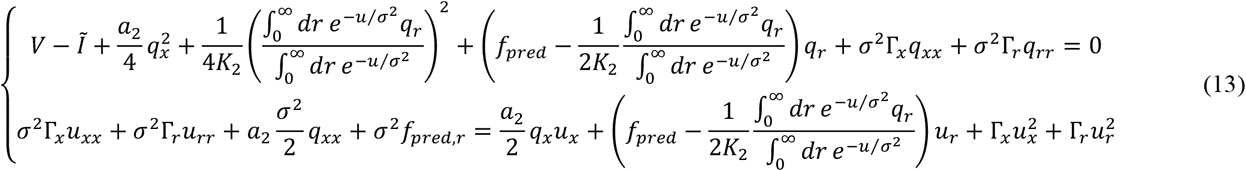

Thus, the original variational problem of minimizing *I* is reduced to system (13) of two integro-differential equations, whose solutions *q*(*x, r*) and *u*(*x, r*) provide the evolutionarily optimized functions *f*_1_(*x, r*), Δ*f*_2_(*x*), and *P*_*stat*_ (*x, r*)

### Series expansions to solve the integro-differential equations

Now, the idea for solving equations (13) is to build series expansions for *u*(*x, r*) and *q*(*x, r*) in terms of *a*_2_ considered as a large parameter (similar to *a* in Paper III^5^) or, alternatively, of 1/*a*_2_ as a small parameter. We think that *a*_2_ is a large parameter because in the limit of *a*_2_ → ±∞ the term in the Lagrangian responsible for maintaining restriction (5) vanishes, which is equivalent to not imposing this restriction, or to the infinitely large value of *M*, which, as we discussed above, agrees with the biophysical interpretation of the problem that *M* is a large (though finite) number. We write the asymptotic behavior of *u*(*x, r*) and *q*(*x, r*) in the following form:

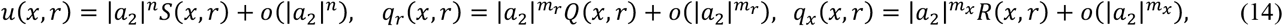

where *n, m*_*r*_ and *m*_*x*_ are constants, and the absolute values of *a*_2_ are used because *n, m*_*r*_ or *m*_*x*_ may be fractional. In general, *m*_*r*_ and *m*_*x*_ may be different. For example, *m*_*r*_ > *m*_*x*_ is possible, if 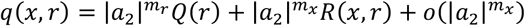, then *q*_*r*_ contains the leading term of the order 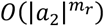, while in *q*_*x*_ this term vanishes.

Some of the combinations of the *n, m*_*r*_ and *m*_*x*_ values can be excluded. For example, if

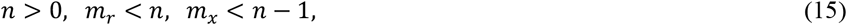

then the leading terms in the second equation in (13) are of the order of *O*(|*a*_2_|^2*n*^), yielding

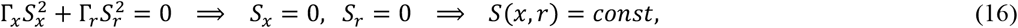

which is not acceptable, because in this case the integral of *P*_*stat*_ (*x, r*) in equation (4) diverges. Also, if

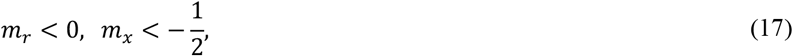

then in the first equation in (13) the leading terms are of the order of 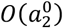, yielding

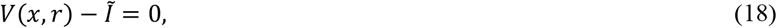

which cannot be true for arbitrary *x* and *r*. Also, if

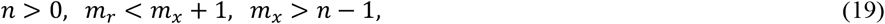

then the only leading term in the second equation in (13) is of the order of 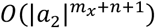, yielding

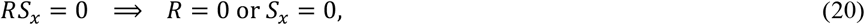

but the first variant is excluded because it contradicts to the assumption that the leading term in *q*_*x*_ is of the order of 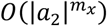, while the second variant is excluded because it leads to a divergence of the integral of *P*_*stat*_(*x, r*) over *x*.

We do not have an analysis of all possible variants of the *n, m*_*r*_ and *m*_*x*_ values yet.

### Particular solution corresponding to the previously considered partial optimization problem

The case considered in Paper III^5^ corresponds to the following values:

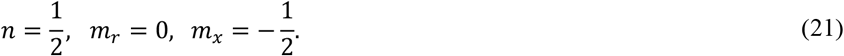

In this case, the leading terms in equations (13) are of the orders of 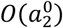 and *O*(|*a*_2_|), respectively:

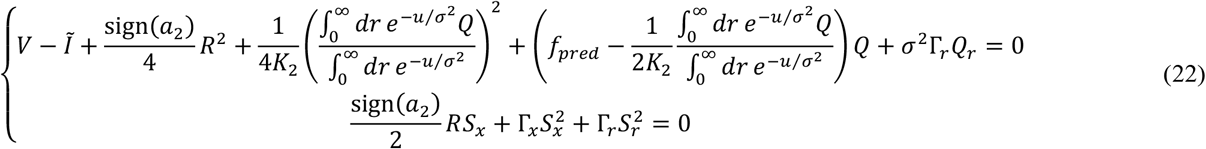

Consider the second of these equations. For brevity and by analogy with equation (9) from paper III,^5^ introduce a notation

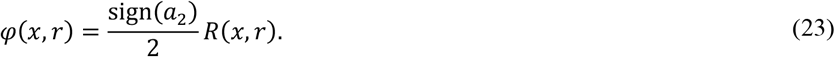

Suppose that for any *r, S*(*x, r*) as a function of *x* has a global minimum at some *x*, which, in general, will depend on *r*: *x* = *x*_*eq*_ (*r*). This assumption is justified by the necessity to ensure convergence of the integral of *P*_*stat*_ (*x, r*) over *x* at any *r*. Then, let us expand *S*(*x, r*) and *φ*(*x, r*) in Taylor series in terms of Δ*x* = *x* − *x*_*eq*_ (*r*) at each *r*:

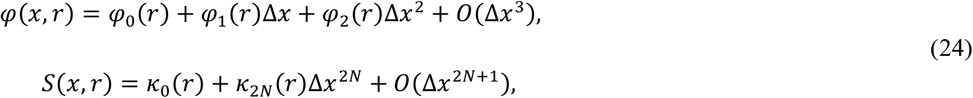

where *N* = 1 or a larger positive integer, and *κ*_2*N*_(*r*) > 0, both in agreement with the assumption about the minimum of *S*(*x, r*). Then, the series for the partial derivatives of *S* are:

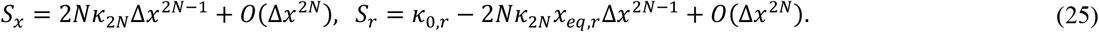

Plugging these expansions into 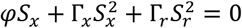, we notice that because *N* ≥ 1, and therefore 2*N* − 1 ≥ 1, the only lowest order term in this equation, namely the *O*(Δ*x*^0^) term, is 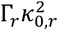. Hence, *κ*_0,*r*_ = 0, so *κ*_0_ does not depend on *r*. Then, the leading terms both in *S*_*x*_ and *S*_*r*_ are of the order of *O*(Δ*x*^2*N*−1^), and *φ* can be found from *S*_*x*_ and *S*_*r*_ as follows:

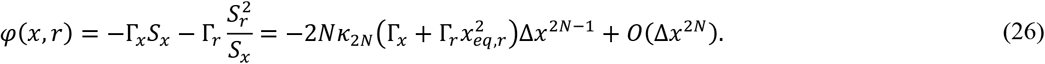

Then, according to equation (23),

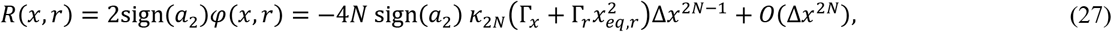

and, according to equations (10) and (14),

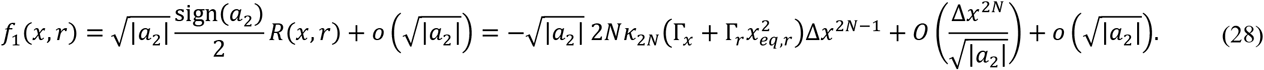

For typical values of Δ*x, u* is finite, that is, *u* ≲ *O*(|*a*_2_|^0^), but at the same time, 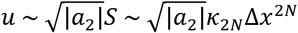, therefore, typical 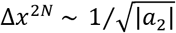, and the term 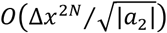 in equation (28) is absorbed into the term 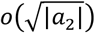.

In the simplest case of *N* = 1, equation (28) simplifies to:

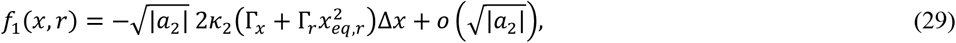

which is essentially the case considered in Paper III,^5^ with

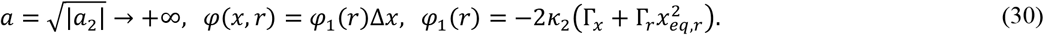

Moreover, if we choose *φ*_1_(*r*) to be constant (a negative constant, because *κ*_2_(*r*) > 0), then the optimal *f*_1_(*x, r*) splits into a sum of two terms, one of which depends only on *x*, while the other depends only on *r*:

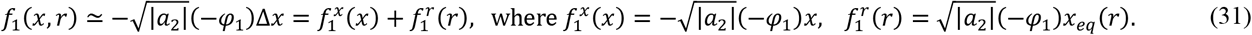

Perhaps we are biased by the known neurobiology, but it should be much easier to implement this separable form of *f*_1_(*x, r*) in specific material mechanisms, because it allows for two separate processes contributing to the temporal dynamics of the state of the neuron, one of which depends only on the current state of the neuron *x* (e.g., based on voltage-gated ion channels), while the other one is purely sensory and depends only on the distance to the predator (e.g., based on ion channels gated by the sensory signal).

With values of *N* > 1, we still retain the key conclusions that the prefactor 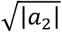 in *f*_1_(*x, r*) goes to infinity, and that at *x* = *x*_*eq*_(*r*), *f*_1_(*x, r*) = 0. What is different is that the terms depending on *x* and terms depending on *r* cannot be separated into two independent contributions anymore, and hence, it may be much more difficult or impossible for the evolution to create material mechanisms implementing this solution for the optimal *f*_1_(*x, r*).

Now consider the first equation in (22). First of all, note that the large *a*_2_ asymptotes for the integrals can be computed, based on the results of the analysis of the first equation. Namely, in the limit of infinitely large *a*_2_, by analogy with equation (24) in Paper III,^5^

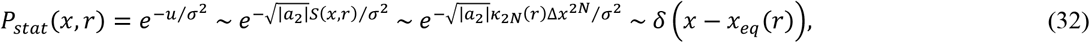

and therefore, the ratio of the integrals has the following asymptotic behavior:

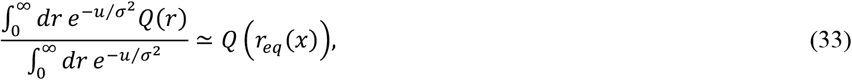

where *r*_*eq*_ (*x*) is the inverse function of *x*_*eq*_ (*r*), that is, ∀*r*: *r*_*eq*_ (*x*_*eq*_ (*r*)) = *r*. Note that in the case under consideration, according to (21), *m*_*r*_ > *m*_*x*_, and therefore, *Q* depends only on *r*, as discussed in the text after equation (14). This emphasizes the necessity of using Δ*f*_2_(*x*), not Δ*f*_2_(*x, r*), in the formulation of the optimization problem (1). With Δ*f*_2_(*x, r*), due to the fact that *Q* depends only on *r*, we would have concluded that Δ*f*_2_ depends only on *r*, and not *x*, which contradicts to the biophysical sense. At the same time, using Δ*f*_2_(*x*), and therefore integration in (11) and (33), allows us to replace this *r* by *r*_*eq*_ (*x*), enabling the dependence of Δ*f*_2_ on *x*.

With equation (33), the first equation in (22) simplifies to:

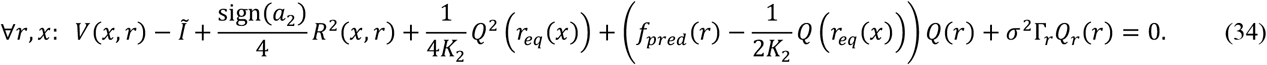

In particular, at *x* = *x*_*eq*_ (*r*)

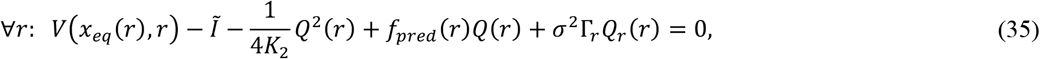

(recall that *R*(*x*_*eq*_(*r*), *r*) = 0, as derived above). Note that equation (35) does not depend on a specific functional form of *R*(*x, r*), and therefore, on the choice of *N* above in the derivation of the optimal *f*_1_(*x, r*).

Note an analogy between this equation and equation (49) in paper III,^5^ and respectively, the analogy *Q* ∼ *U*_*r*_. Based on it, we make the following substitution:

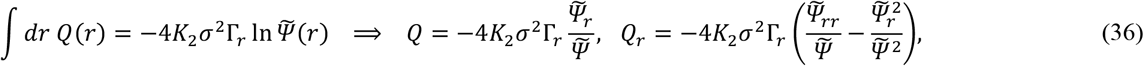

and then differential equation (35) transforms to

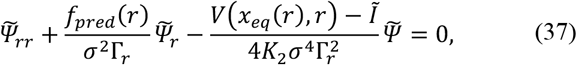

which is similar to, but does not exactly coinside with equation (52) from paper III.^5^ The similarity is that both are second-order differential equations, with the prefactor for the first order derivative including *f*_*pred*_(*r*) and with the linear term proportional to *V* minus the correspondign Lagrange multiplier. The differences are: (1) the prefactor for the first order derivative in equation (52) from paper III^5^ includes also terms depending on *φ*_1_, *φ*_2_ and *x*_*eq*_ ; (2) the constant coefficients before *f*_*pred*_ differ by a factor of 2, and the constant coefficients before *V* − *Ĩ* differ by a factor of 4; (3)the optimal function Δ*f*_2_(*x*) is expressed in terms of *Ψ* and 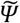with constant prefactors that differ by a factor of 2, namely,

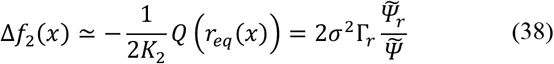

in this work, versus

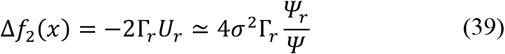

in Paper III.^5^ The reasons for these differences between the solution to the general problem (1) as given in this work, and the solution to the particular problem as given in Paper III,^5^ are not clear to us.

Now, consider the asymptotic behavior of the solutions at small and large *r*. In the limit of *r* → +∞, assuming that *f*_*pred*_ =*σ*^2^Γ_*r*_ /*r* − *εr* as above, we get the differential equation

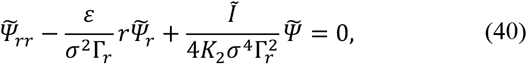

which has the following asymptotic solution:

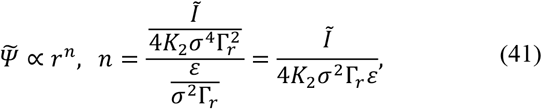

which, taking into account equation (38), corresponds to the following Δ*f*_2_(*x*):

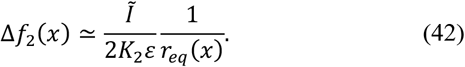

This result is the same as in equation (57) derived earlier in Paper III^5^ for the partial optimization problem (assuming that the Lagrange multiplier *Ĩ* equals the minimized value of the functional *I*). As before, in the limit of *r*_*eq*_ (*x*) → +∞, Δ*f*_2_ vanishes: Δ*f*_2_ → 0^+^.

The asymptotic behavior of the differential equation (37) at *r* → 0^+^ is:

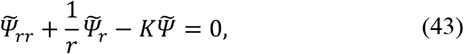

where

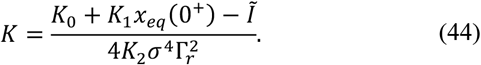

Therefore, the asymptotic behavior of the solution at *r* → 0^+^ is

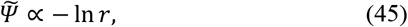

which, taking into account equation (38), corresponds to the following Δ*f*_2_(*x*):

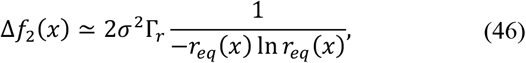

which is different from the corresponding asymptotic (64) from paper III^5^ derived for the partial optimization problem. However, in the limit of *r*_*eq*_ (*x*) → 0^+^, it is still true that Δ*f*_2_ → +∞, as before.

### Possible stepwise character of the optimal motor function Δ*f*_2_

Finally, note that this derivation also predicts, under certain conditions, a stepwise increase in Δ*f*_2_(*x*) at the value of *x* that corresponds to *r*_*d*_, the distance at which *V*(*x, r*) undergoes a pointwise decrease, according to equation (2). Note that in the limit of *r* → +∞, for the asymptotic solution (41), the leading terms in the differential equation are the first-order derivative and linear terms, while the second-order derivative is of lower order in *r*. If this relationship still holds in the vicinity of *r*_*d*_, then the original equation (37) simplifies to

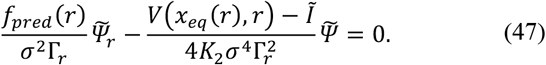

Hence, a discontinuous change in *V* by Δ*V* = *V*(*x*_*eq*_(*r*_*d*_ + 0^+^), *r*_*d*_ + 0^+^) − *V*(*x*_*eq*_ (*r*_*d*_ − 0^+^), *r*_*d*_ − 0^+^) = −*K*_0_ causes a discontinuous change in 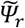by

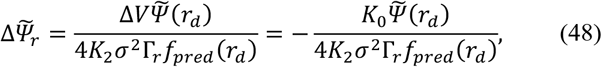

and therefore, a stepwise increase in Δ*f*_2_ by ΔΔ*f*_2_ that can be found equations (38) and (48):

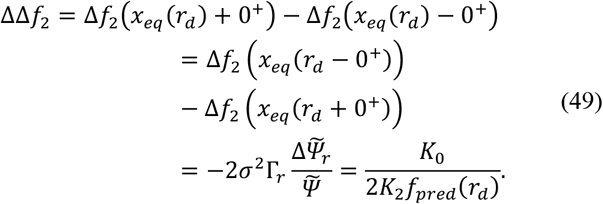

This derivation is valid, however, only if the second-order term in the original equation (37) is small at *r* = *r*_*d*_. Otherwise, the discontinuous change in *V* by Δ*V* will lead to a discontinous change in 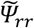, not 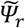, so the increase in Δ*f*_2_ will be continous, and the effect of Δ*V* will spread to a finite interval of *r*.

## Discussion

In this work, we present a solution to the general optimization problem. It leads to qualitatively (and sometimes even quantitatively) similar key results to those obtained in Paper III^5^ for the partial optimization problem. For example, the deterministic (non-noisy) part of the temporal derivative of the membrane potential of a neuron is proportional to the deviation of the current value of the membrane potential from its stationary value that depends on the sensory input. The motor reaction depends on the neuronal potential in such a way that when the neuron is not excited, the motor reaction is absent, while neuronal excitation triggers a motor reaction.

However, the general problem presented here is broader. The solution analogous to that from Paper III^5^ is obtained for a specific choice of parameters *n, m*_*r*_ and *m*_*x*_ given by equation (21).

Alternative solutions for other values of *n, m*_*r*_ and *m*_*x*_ might formally exist and might even have a reasonable biophysical interpretation, alternative to the one implied by the original motivation of this simplest model. This problem of such alternative solutions of the general optimization problem remains open.

In the previous papers,^1,2,5^ we focused on the statement that the minimized functional *I* can be made arbitrarily close to zero by taking certain limits for parameters from *f*_1_(*x, r*) and Δ*f*_2_(*x*), such as *a* → +∞, λ → +∞, etc., in other words, the evolutionary cost of the nervous system can be made arbitrarily small. However, these infinite solutions for *f*_1_(*x, r*) and/or Δ*f*_2_(*x*) are not realistically achievable, given possible material mechanisms implementing a nervous system (e.g., ion channels). In this work, for the first time, we demonstrate that the restriction of finite speeds of neuromolecular processes, given by equation (5), leads to possibly large but still finite values of parameters *a* or |*a*_2_|, and hence, finite solutions for *f*_1_(*x, r*) and Δ*f*_2_(*x*) [see equations (31) and (38)]. From the viewpoint of neurobiology, this means that optimal membrane voltage kinetics, optimal response functions of sensory neurons, and optimal effector (motor) response functions exist and are finite. In particular, high – but not infinitely high – permeability and sensitivity of ion channels, as measured by *M* in equation (5), leads to sharp (narrow and high) – but not infinitely sharp – neuronal spikes as the evolutionary optimal solution. The specific shapes of such spikes (not just numerical parameters in pre-set functions, but functions themselves) can be obtained as a solution to the variational problem.

Some questions still remain open. In particular, alternative solutions to the general problem may exist, as mentioned above. If such solutions exist for *n* ≤ 0, then the probability distribution *P*_*stat*_ (*x, r*) does not tend in such cases to the delta function in the limit of *a* → +∞ or *M* → +∞, resulting in a very different biophysical interpretation of the solution, including the absence of sharp neuronal spikes. Whether such solutions exist is not known to us. Also, is it true in general that every optimal *f*_1_(*x, r*) is such that the equation *f*_1_(*x, r*) = 0 at each *r* has only one solution *x* =*x*_*eq*_ (*r*)? Or do multiple stationary states of a neuron at certain *r* exist? On the technical side, it is not clear to us why the optimal functions Δ*f*_2_(*x*), derived in this work for the general solution and in the previous work for the particular solution [see equations (37), (38), (39) in this work, and equation (52) from paper III^5^], are somewhat different from each other. All these questions require further investigation.

How different are the general solutions presented here, independent of specific molecular mechanisms and material implementations of nervous systems, from the properties of real simple nervous systems? This question requires further empirical investigations. We expect that over hundreds of millions of years that simple nervous systems exist, evolutionary optimization of their molecular mechanisms has nearly exhausted all possible optimization, and therefore, the real properties of nervous systems (membrane voltage kinetics, sensory and effector response functions) are close to those predicted by the presented theory, given that the optimization is performed for the appropriate expression for the cost of the work of a nervous system.

In conclusion, this work demonstrates that well-known basic properties of the nervous system, such as sharp neuronal spikes, or response functions of sensory neurons and effectors, can be strictly derived from the fundamental principles of evolutionary optimization.

